# Transportome remodeling of a symbiotic microalga inside a planktonic host

**DOI:** 10.1101/2024.06.01.596945

**Authors:** C Juery, A Auladell, Z Füssy, F Chevalier, DP Yee, E Pelletier, E Corre, AE Allen, DJ Richter, J Decelle

## Abstract

Metabolic exchange is one of the foundations of symbiotic associations between organisms and is a driving force in evolution. In the ocean, photosymbiosis between heterotrophic host and microalgae is powered by photosynthesis and relies on the transfer of organic carbon to the host (e.g. sugars). Yet, the identity of transferred carbohydrates as well as the molecular mechanisms that drive this exchange remain largely unknown, especially in unicellular photosymbioses that are widespread in the open ocean. Combining genomics, single-holobiont transcriptomics and environmental metatranscriptomics, we revealed the transportome of the marine microalga *Phaeocystis* in symbiosis within acantharia, with a focus on sugar transporters. At the genomic level, the sugar transportome of *Phaeocystis* is comparable to non-symbiotic haptophytes. By contrast, we found significant remodeling of the expression of the transportome in symbiotic microalgae compared to the free-living stage. More particularly, 32% of sugar transporter genes were differentially expressed. Several of them, such as GLUTs, TPTs and aquaporins, with glucose, triose-phosphate sugars and glycerol as potential substrates, were upregulated at the holobiont and community level. We also showed that algal sugar transporter genes exhibit distinct temporal expression patterns during the day. This reprogrammed transportome indicates that symbiosis has a major impact on sugar fluxes within and outside the algal cell, and highlights the complexity and the dynamics of metabolic exchanges between partners. This study improves our understanding of the molecular players of the metabolic connectivity underlying the ecological success of planktonic photosymbiosis and paves the way for more studies on transporters across photosymbiotic models.

## INTRODUCTION

Symbiosis between a heterotrophic host and a photosynthetic partner (photosymbiosis) is considered to be the primary event which led to the acquisition and distribution of plastids in the evolution of eukaryotes [1, 2]. Photosymbiosis remains an essential life strategy and supports the functioning of today’s aquatic ecosystems especially in oligotrophic waters [3–6]. This partnership, considered as mutualistic in the spectrum of symbioses, can provide a competitive advantage in a nutritionally challenging environment where nutrients and prey are scarce (oligotrophic waters). Therefore, partners need to establish a metabolic connection in order to exchange metabolites and nutrients. The microalgae need to be supplied by the host with all the essential macro- and micro-nutrients (e.g. iron, nitrogen) to maintain their metabolic and physiological activity. In turn, the host can benefit from photosynthetic products (photosynthates) exported from the microalgae [7]. In the past decade, nanoSIMS (Nanoscale Secondary Ion Mass Spectrometry) studies coupled with ^13^C labeling improved our knowledge on carbon transfer and allocation in different photosymbiotic system [8–10]. Transferred photosynthates have been mostly investigated in benthic multicellular photosymbioses such as reef-dwelling invertebrates (e.g. anemones, jellyfish, giant clams) living with Symbiodiniaceae microalgae. Sugars, which are the main photosynthetic products and lipids are considered to be the main photosynthates exported from the symbiotic microalgae [11–13]. Glucose was shown to be a major transferred metabolite in some photosymbioses [14, 15], as well as inositol, galactose, and galactosylglycerol [16, 17]. Glycerol was also suggested as a putative transferred metabolite since it is significantly released by free-living Symbiodiniaceae in culture [18–20]. However, the exact nature of translocated carbohydrates is still uncertain since experimental evidence is difficult to obtain on such photosymbiotic systems and on these very rapid metabolic processes.

In symbiosis, both partners need to reprogram their transportome (defined as “membrane proteins responsible of the translocation of any kind of solutes across the lipid layer” [21]) in order to establish metabolic connectivity. Most metabolites including sugars require a complete set of transporters to traverse algal and host membranes. For example, in symbioses between plants and fungi, changes in expression of the transporter genes are essential to connect and integrate different metabolisms [22]. In marine photosymbioses, some transporters have been highlighted in genomic [23] and transcriptomic studies [24]. A glucose transporter (GLUT8) and an aquaporin (GflP) that could putatively transport glycerol, were described in anemones and the jellyfish *Cassiopea* [25, 26]. While most studies have focused on host transporters, less is known about the ones of the symbiotic microalgae that can export energy-bearing metabolites derived from photosynthesis. So far, a SWEET (Sugars Will Eventually be Exported Transporter) has been described as a glucose transporter located in the cell membrane of the microalga *Brevolium* (Symbiodiniaceae), symbiont of the anemone *Exaiptasia diaphana* [27].

Relatively less studied than reef ecosystems, a wide diversity of photosymbioses are also found in marine and freshwater plankton. For instance, radiolarians and foraminiferans that are widespread in the sunlit layer of the ocean can host diverse microalgae [28]. Among radiolarians, some species of acantharia live in symbiosis with the microalga *Phaeocystis* (Haptophyta) in different oceanic regions [29]. Symbiotic acantharia significantly contribute to the primary production (up to 20% in surface oligotrophic oceans) and carbon fluxes to deep layers of the ocean [30, 31]. Previous studies using 3D electron microscopy have shown that the microalga *Phaeocystis* undergoes drastic morpho-physiological changes in symbiosis: cell and plastid volume, as well as plastids number, greatly increased compared to free-living cells in culture [32]. In addition, while cell division is very likely arrested in symbiosis, photosynthesis and carbon fixation are enhanced, corroborated by an upregulation of many genes of the Calvin-Benson cycle. Symbiotic microalgae with their expanded photosynthetic apparatus therefore produce a significant amount of organic carbon but the identity of these compounds and the mechanisms by which they are transferred to the host remain largely unknown. Investigating the composition and expression of the algal transportome can reveal how symbiotic microalgae metabolically connect to its host and can provide insights on the putative exchanged metabolites.

Here, we conducted genomic and transcriptomic analyses on an uncultivable planktonic photosymbiosis between the microalga *Phaeocystis* and acantharian hosts in order to shed the light on the molecular mechanisms of their metabolic connectivity. More specifically, we investigated whether the algal transportome is remodeled in symbiosis. We first compared sugar transporter genes in haptophyte genomes and studied their expression in free-living and symbiotic stages of *Phaeocystis* using a combination of single-holobiont transcriptomics and *in situ* environmental metatranscriptomics. We evaluated the transcriptional dynamics of these sugar transporters in symbiosis at different periods of the day. This study reveals that the transportome of the microalga is significantly remodeled in symbiosis within a host and pinpoints putative key sugar transporters with different transcriptional patterns during the day. This work significantly improves our understanding of the metabolic connectivity between a host and microalgae and so provides fundamental knowledge of the ecological success of this widespread symbiosis in the ocean.

## RESULTS AND DISCUSSION

### Genomic inventory of sugar transporters in *Phaeocystis* species

To understand the molecular toolbox underlying metabolic fluxes, we first unveiled the transportome of the microalga *Phaeocystis* at the genomic level. We determined the different categories of transporters and their proportions among six haptophytes species in order to reveal a specific genomic footprint in *Phaeocystis* potentially linked to its symbiotic lifestyle, compared to non-symbiotic related haptophytes (Fig. 1A). We also predicted subcellular localization of transporters using a combination of different *in silico* tools. We particularly focused on sugar transporters, since soluble sugars can be the main exported currency to the host. Using the same method of protein annotation for each species, we identified transporters based on the presence of transmembrane domains and protein domain annotations using the InterPro/Pfam classification. In total, we found 270 unique Pfam domains for the transporter genes in haptophytes genomes. The three different *Phaeocystis* species analyzed here contained an average of 965 transporter genes corresponding to 3% of all genes (1,244 genes or 4% if we include those lacking a predicted transmembrane domain, see Methods, Fig. 1A, Table S1, “General values1” and “General values2 (TMD)”). By comparison, 2 and 3.5% of transporter genes were found in the genomes of two symbiotic microalgae *Symbiodinium microadriaticum* and *Brevolium minutum*, respectively (based on Joint Genome Institute’s genome portal annotations).

**Fig. 1.**
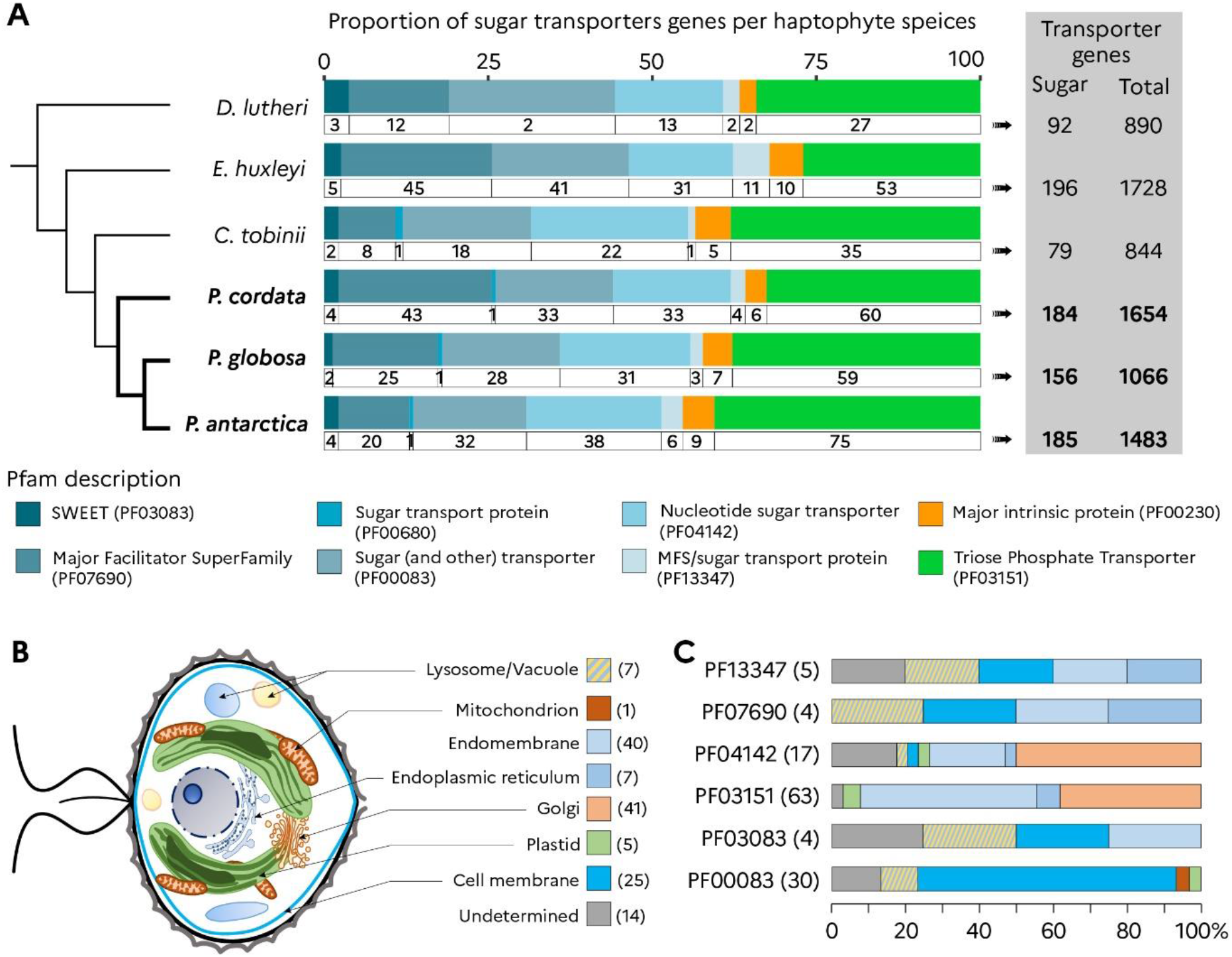
Genomic composition of sugar transporters of six haptophyte species. **A)** Gene composition and Pfam families of different sugar transporters found in the genomes of six haptophytes species represented in a schematic phylogenetic tree (*Diacronema lutheri, Emiliania huxleyi, Chrysochromulina tobinii, Phaeocystis antarctica, Phaeocystis globosa, Phaeocystis cordata*). Eight Pfam families were found and represented by different colors and the number of sugar transporter genes within each Pfam is indicated. The genus *Phaeocystis* was found to contain between 156 and 184 sugar transporters out of 1066 to 1654 transporters in total. **B)** In *Phaeocystis cordata,* the subcellular localization and number of sugars transporters are shown in the schematic drawing of the microalga (prediction using a combination of *in silico* tools, see methods and Table S1). **C)** For each Pfam, the proportion of sugar transporters within each compartment of the cell are shown in the bar chart.

Across *Phaeocystis* species genomes, sugar transporter genes represented on average 18% (175 out of 965) of all transporter genes and were classified into eight different core Pfams (i.e. shared among all haptophyte species, Fig. 1A, Table S1). The largest Pfam family of sugar transporters is the Triose Phosphate Transporters (TPT, PF03151) with 64.7 genes on average across *Phaeocystis* species, similar to *Emiliania huxleyi*, but higher than in diatom genomes (between 13 and 22 TPTs for *Phaeodactylum tricornutum* [33, 34]). In plants, TPTs are transporters located in the plastid envelope and export photosynthetically-derived sugars to the cytosol [35]. For *P. cordata*, *in silico* subcellular localization analyses predicted only three out of 60 TPTs associated to the plastid membranes while the majority was predicted to be located in the endomembrane system (Golgi apparatus or endoplasmic reticulum, ER) (Fig. 1C Table S1). Similar to diatoms, the localization of TPTs in the four membranes of the secondary red plastid of *Phaeocystis* remains ambiguous and TPTs could also be localized elsewhere in the cell [36–38].

The second largest sugar transporter family was the “Sugar and Other transporters” Pfam PF00083, containing 31 genes in *Phaeocystis* on average (41 in *Emiliania huxleyi*, Fig. 1A). This Pfam is composed of different transporters with various substrates, such as glucose, galactose, mannose, polyol, and inositol sugars [39, 40]. For instance, using a complementary Hidden Markov Model (HMM) search of InterPro domains in *P. cordata* PF00083 proteins, we found 5 GLUT transporter (IPR002439), 6 Sugar transporter ERD6/Tret1-like (IPR044775), and 12 Sugar transport protein STP/Polyol transporter PLT (IPR045262) domains (Table S1, “GLUTcharacterization”). *In silico* subcellular localization successfully assigned a prediction for 21 of the 33 transporter genes from PF00083 of *P. cordata* in the cell membrane, three in vacuoles, and one in the plastid membrane (Fig. 1B). These results suggest that a large part of these transporters might be involved in sugar flux at the cell surface. Nucleotide sugar transporters (NSTs, PF04142) play an important role in the biosynthesis of glycoproteins, glycolipids and non-cellulosic polysaccharides translocating nucleotide-sugars in the Golgi [41, 42]. In *Phaeocystis* species, we found 34 NSTs genes on average (31 in *E. huxleyi*). In *P. cordata*, 50% of the NSTs were predicted to be localized in the Golgi apparatus, in accordance with their known biological function (Fig. 1C).

SWEET transporters (Sugars Will Eventually be Exported Transporters, PF03083) are bidirectional transporters of small sugars following the concentration gradient [43]. SWEETs have been highlighted in terrestrial and aquatic symbioses (Fabaceae- *Rhizobium*, cnidarians-Symbiodiniaceae), particularly in sugar efflux from the photosynthetic to the heterotrophic partner [44–47]. Between two and four SWEET genes were found across *Phaeocystis* species genomes. By comparison, many duplications and diversification of SWEET genes are known in plant genomes (17 in *Arabidopsis thaliana* and up to 53 in *Glycine max* [48]). Among the four SWEET proteins of *P. cordata*, one was predicted in the lysosome/vacuole, one in the endomembrane system, and one in the cell membrane. For the Pfam PF13347 (MFS/sugar transport protein) [49, 50], we found four genes in *Phaeocystis* genomes In addition, we investigated the presence of putative sugar transporters from the Major Facilitator Superfamily PF07690 (MFS). Through an HMM search (Table S1, “PF07690_characterization”), we detected 43 putative sugar transporters for *P. cordata* and 25 and 20 for *P. globosa* and *P. antarctica*, respectively. Among them, many genes corresponded to glucose-6-phosphate transporter (SLC37A1/SLC37A2,IPR044740), which are on the chloroplast membrane in plants [51]. In *Phaeocystis*, most of these PF07690 transporters were predicted to be either at the cell membrane or ER (Table S1, “Subcellular Localization P.cord”). Finally, we investigated the presence of genes belonging to PF00230 that corresponds to a specific type of aquaporin, found to be involved in different reef photosymbioses for putative glycerol transport (anemones, jellyfish, and giant clams [25, 26, 52]). In *Phaeocystis* genomes, seven (*P. cordata*) to nine (*P. antarctica*) homologs of these aquaporin genes were found.

This inventory of sugar transporters in genomes unveiled different categories that could be involved in the influx and efflux of sugars in the microalga *Phaeocystis*. Overall, 25 sugar transporters were predicted to be localized at the cell membrane (Fig. 1B), and the vast majority presented a putative localization in the endomembrane system (including ER or Golgi). We also hypothesize that sugar transporters can be located on vesicles derived from endomembrane system that could fuse to the cell membrane [53]. We did not find any evidence of large copy number variations of sugar transporters genes in *Phaeocystis* genomes compared to other haptophytes, which could have been a genomic footprint to explain the predominance of this genus in symbiosis. For instance, it has been demonstrated that symbiotic *Symbiodinium* clades presented enriched functions related to transmembrane transport in their genomes, especially for the major facilitator superfamily (PF07690) [54]. This genomic characterization generates fundamental knowledge on this key marine phytoplankton taxon and is an essential step for unveiling the expression dynamics of sugar transporter genes in symbiosis.

### The algal transportome is significantly remodeled in symbiosis

To reveal which sugar transporters might play a role in symbiosis, we assessed their gene expression based on single-holobiont transcriptomic analyses. More specifically, we compared the expression of transporter genes of the microalga *Phaeocystis cordata* between free-living (four and five culture replicates in exponential and stationary growth phases, respectively) and symbiotic conditions (17 holobionts representing five host species collected in the Mediterranean Sea) through a Differential Expression (DE) analysis (Fig. 2 and S1). Each sample was frozen at the same period of the day (late evening, 7pm). About 75 million of reads were obtained per holobiont sample, producing a total of 1,950 billion of reads in this study. A *de novo* reference transcriptome from total RNA sequences of cultured *P. cordata* was built and used to quantify gene expression.

**Fig. 2.**
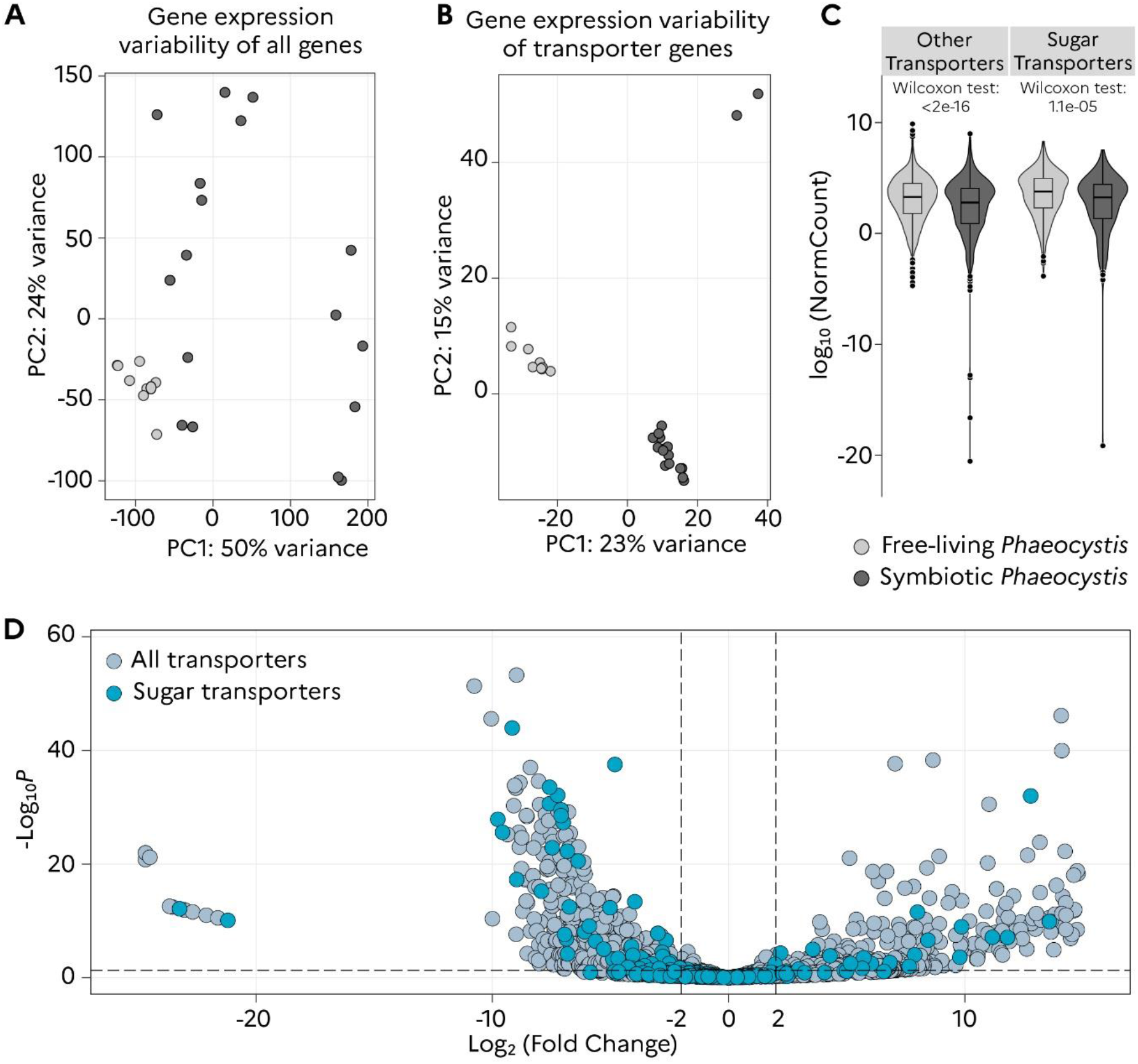
Expression of algal transporter genes between the free-living and symbiotic stages of the microalga *Phaeocystis cordata*. **A)** Principal Component Analysis (PCA) of gene expression variance between free-living (culture; light gray) and symbiotic *Phaeocystis cordata* (dark gray), including all the expressed genes; X-axis represent variance according to sample types and y-axis according to replicates within each sample type. **B)** PCA of expression variance of the 2,133 transporter genes in the free-living and symbiotic phases of *Phaeocystis cordata*. **C)** Comparison of normalized read count values of the sugar and all others transporter genes between the free-living and symbiotic stages of *Phaeocystis cordata* (Wilcoxon rank sum test). **D)** Volcano plot showing the fold change of up- and down-regulated transporter genes of symbiotic *Phaeocystis*, compared to the free-living stage, including sugar transporters (blue circles) and other transporters (grey circles) following DESeq2 analysis (log2FC |2|, p-adj <0.05, see Table S3).

Before comparing the expression of the transportome between the free-living and symbiotic stages, we assessed whether the transportome of free-living *Phaeocystis* cells maintained in culture varies with respect to the growth phase (i.e. exponential *vs* stationary phase). Only 3.3% of the transporters were found to be differentially expressed between exponential and stationary phase, indicating that the expression of the transportome is not drastically modified (Table S2). We therefore considered hereafter both growth phases as free-living replicates to compare with the symbiotic stage.

From the 2133 transporters genes considered as expressed in this analysis (sum of normalized read counts > 10 in all replicates, Table S3), 52% (1115 genes) were found to be expressed exclusively in free-living and 12% (263 genes) exclusively expressed in symbiosis (Table S3, “Exp FL Symb”). A principal component analysis (PCA) of the DESeq2 dataset showing the gene expression variance revealed: i) a clear separation between symbiosis and free-living samples; ii) a high variance in expression of all genes across symbiotic replicates, and iii) clustering of expression of transporter genes in symbiotic samples (even if a lower part of the variance −38%- is explained by the first two dimensions of the PCA for transporter genes compared to all genes −74%-, Fig. 2A and 2B). We verified this clustering pattern by using different subsets of expressed genes with similar numbers of genes as controls (Fig. S1). These results show the existence of two distinct transportomes of the microalga *Phaeocystis* expressed in the symbiotic and free-living stages.

Compared to the free-living stage, 42% of the transportome was significantly remodeled in symbiosis with 888 differentially expressed genes (Fig 2C). Among those, 26% (556 genes) were downregulated and 16% (332 genes) were upregulated in symbiosis (Table S3, “UP-DOWN global”). These numbers are higher than the ones found in the symbiotic *Brevolium,* and with an opposite trend: 213 up-regulated and 167 down-regulated [24]. We explain this remodeling of *Phaeocystis* transportome in symbiosis by a significant global decrease of the transporter gene expression (normalized read counts, Wilcoxon rank sum test, p-value <0.01, Fig. 2C) and high positive fold changes for some transporter genes (Fig 2D, aquaporin PF00230 with log2FC = 13.58, Table S3). Note that 58% of the algal transportome remained expressed in symbiosis but without differential expression. Therefore, these results suggest that many algal transporters could be less required in symbiosis likely due to the transition from the ocean to the host microhabitat, but some transporters could be specifically induced in symbiosis with high transcriptional activity in response to metabolic changes within a host and potentially enable the metabolic connectivity between the two partners.

### Expression of the algal sugar transporters in symbiosis

We next focused on the expression of sugar transporter genes of the microalga in symbiosis. From our dataset, 223 genes were found to be expressed (out of the 330 sugar transporter genes in the reference transcriptome; TableS3 “UP-DOWN global“ and “*P.cordata* SugarTR Anno”). At the Pfam level, no significant changes of the global gene expression of a family were observed between free-living and symbiotic microalgae (Fig. 3A). Yet, at the gene level, 32% of sugar transporters were remodeled in symbiosis with 9 % (19 genes) upregulated and 23% (52 genes) downregulated (Fig. 2D, Table S3, “UP-DOWN global”). Note that 68% (152 genes out of the 223) of the sugar transporter genes were still expressed in symbiosis (named “neutral” in Table 1). Among the 19 up-regulated genes in symbiosis, we found ten TPTs (Triose Phosphate Transporters, PF03151, average log2FC = 6.4), two Aquaporins (PF00230, average log2FC = 14.4), two monosaccharide transporters (PF00083, average log2FC = 4.3), one MFS/sugar transport protein (PF13347, average log2FC =5.7), two Nucleotide sugar transporters (PF04142, average log2FC = 5.4) and two genes from PF07690 (annotated as Glycerol-3-P transporter, average log2FC = 4.1) (Fig. 3B, Table 1). These transporters therefore participate to the significant remodeling and specialization of the algal sugar transportome in symbiosis and potentially play a key role in the flux and exchange of sugars.

**Fig. 3.**
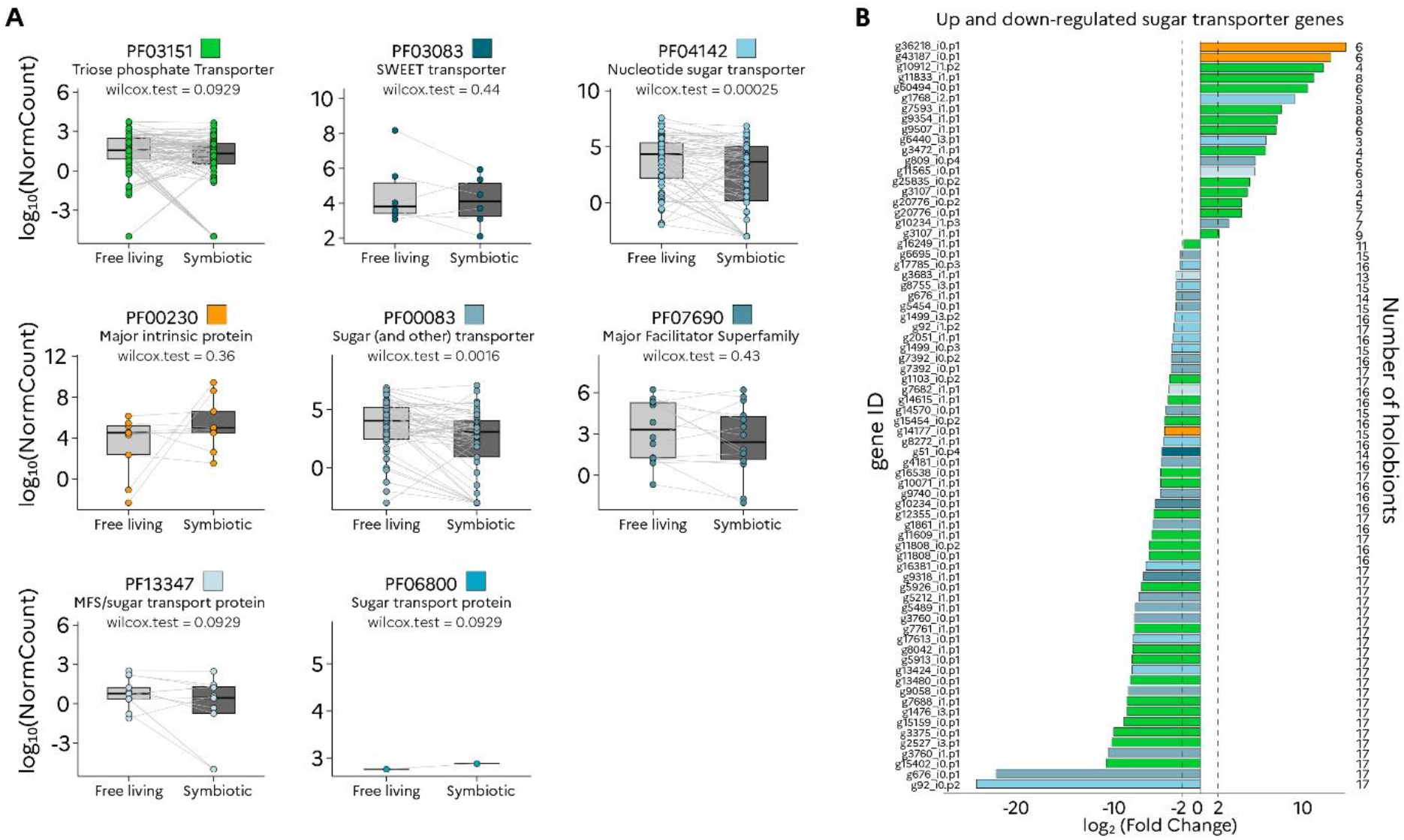
Expression of sugar transporter genes between the free-living and symbiotic *Phaeocystis cordata* at the gene and Pfam level. **A)** Pairwise comparison of the normalized read counts of sugar transporter genes from eight Pfams (shown by different colors) between free-living and symbiotic *Phaeocystis cordata* (Wilcoxon rank sum test, sum of read counts >10 over all replicates). Each dot and line represents one sugar transporter gene. **B)** Distribution of fold changes for up- and down-regulated sugar transporter genes in symbiotic *Phaeocystis* compared to free-living (log2FC > 2, *p*-adj < 0.05). Each line corresponds to one sugar transporter gene from a specific Pfam, which was found to be up- or down-regulated in a different number of holobionts (Y axis, right, see also Table S3).

**Table 1.** Differential expression of sugar transporter genes from different Pfams in the symbiotic stage of *Phaeocystis cordata* compared to the free- living one. (log2FC |2|, *p*-adj < 0.05). Note that ”neutral” indicates that the sugar transporter gene is expressed in symbiosis, but not significantly down- or up- regulated compared to the free-living stage.

|  | Annotation |  | -2< log2FC< 2 - padj 0.05 |  |  |  |
| --- | --- | --- | --- | --- | --- | --- |
|  | Pfam | Pfam_description | Down | Neutral | UP | Total |
| Sugar Transporters | PF00083 | Sugar_(and_other)_transporter | 13 | 28 | 2 | 43 |
|  | PF03083 | Sugar_efflux_transporter_for_intercellular_exchange | 1 | 5 | 0 | 6 |
|  | PF03151 | Triose-phosphate_Transporter_family | 19 | 56 | 10 | 85 |
|  | PF04142 | Nucleotide-sugar_transporter | 10 | 30 | 2 | 39 |
|  | PF07690 | Major_Facilitator_Superfamily | 2 | 9 | 2 | 13 |
|  | PF00230 | Major_Intrinsic_protein | 1 | 6 | 2 | 13 |
|  | PF13347 | MFS/sugar_transport_protein | 2 | 5 | 1 | 7 |
|  | PF00474 | Sodium/glucose_cotransporter | 0 | 1 | 0 | 1 |
|  | PF00892 | GDP-fucose_transporter | 0 | 1 | 0 | 1 |
|  | PF06800 | Sugar_transport_protein | 0 | 1 | 0 | 1 |
|  | PF08489 | UDPxylose / SugarPhosphate translocator | 2 | 0 | 0 | 2 |
|  | - | L-arabinose | 1 | 0 | 0 | 1 |
|  | - | GDP-fucose | 1 | 6 | 0 | 7 |
|  | - | GDP-mannose | 0 | 4 | 0 | 4 |
| Total |  |  | 52 | 152 | 19 | 223 |

Concerning the TPTs, only 10 out of 85 genes (12%) were upregulated and 66% were still expressed in symbiosis. It is possible that the observed upregulation of some TPTs can be linked to the multiplication of plastids in symbiosis (from two in free-living to up to 60 plastids in symbiosis [32]) and would ensure enhanced sugar export into the cytosol. The most upregulated sugar transporter genes correspond to two aquaporins (PF00230, log2FC of 15.1 and 13.6) (Fig. 3B, Table 1, Table S3). This type of aquaporin, also known as GlpF (Glycerol Facilitator), was found to be involved in two benthic photosymbioses and suggested to be involved in glycerol transport [25, 26]. The four upregulated transporter genes with glycerol as putative substrate (two genes of PF00230 and two genes of PF07690) raise the hypothesis that this metabolite can be important for the carbon metabolism of the holobiont and possibly transported to the host. Two other upregulated genes corresponded to putative transporters of monosaccharides (PF00083), with assignment to GLUT proteins (Fig. S2 and Table S1, “GLUT Characterization”). One of these two GLUT proteins was predicted to be localized at the cell membrane (Table S3, “SubcellLoc SugarTRUP”). Four SWEET transporter genes (PF03083) were expressed in symbiosis (log2FC ranging between −1.5 to 0.90), but not upregulated. The upregulated MFS/sugar transport protein (PF13347, log2FC = 5.7) could also play a role in sugar flux as shown in plants [50]. This transcriptomic analysis from freshly collected holobionts allowed us to reveal the sugar transportome expressed in the symbiotic microalga and identify candidate genes for future functional characterization. To have an alternative line of evidence, we investigated the expression of this algal transportome *in situ*, exploiting metatranscriptomic data collected in the Mediterranean Sea.

### *In situ* sugar transportome expression of the microalga *Phaeocystis* in the Mediterranean Sea

Using the *Tara* metatranscriptomic dataset from the Mediterranean Sea, we evaluated the expression of sugar transporter genes of the microalga *Phaeocystis* in two size fractions (small: 0.8–5 µm; and large: 180–2000 µm) collected in surface waters of six stations from the *Tara* Oceans expedition ([55], Fig. 4A). The small size fraction mainly corresponds to the free-living stage of *Phaeocystis cordata* (4 µm in size [56]) while its symbiotic stage within acantharians is mainly detected in the large size fraction (note that the Mediterranean species, *Phaeocystis cordata,* does not form colonies [57]). From this metatranscriptomic dataset, we identified 270 sugar transporters with a difference between the large (123 genes) and small (266 genes) size fraction. This can be partially explained by the lower abundance of *Phaeocystis* transcripts in the 180-2000 µm fraction that might have been diluted and so less sequenced due to the high abundance of transcripts from large multicellular organisms (zooplankton) (Fig. 4B).

**Fig. 4.**
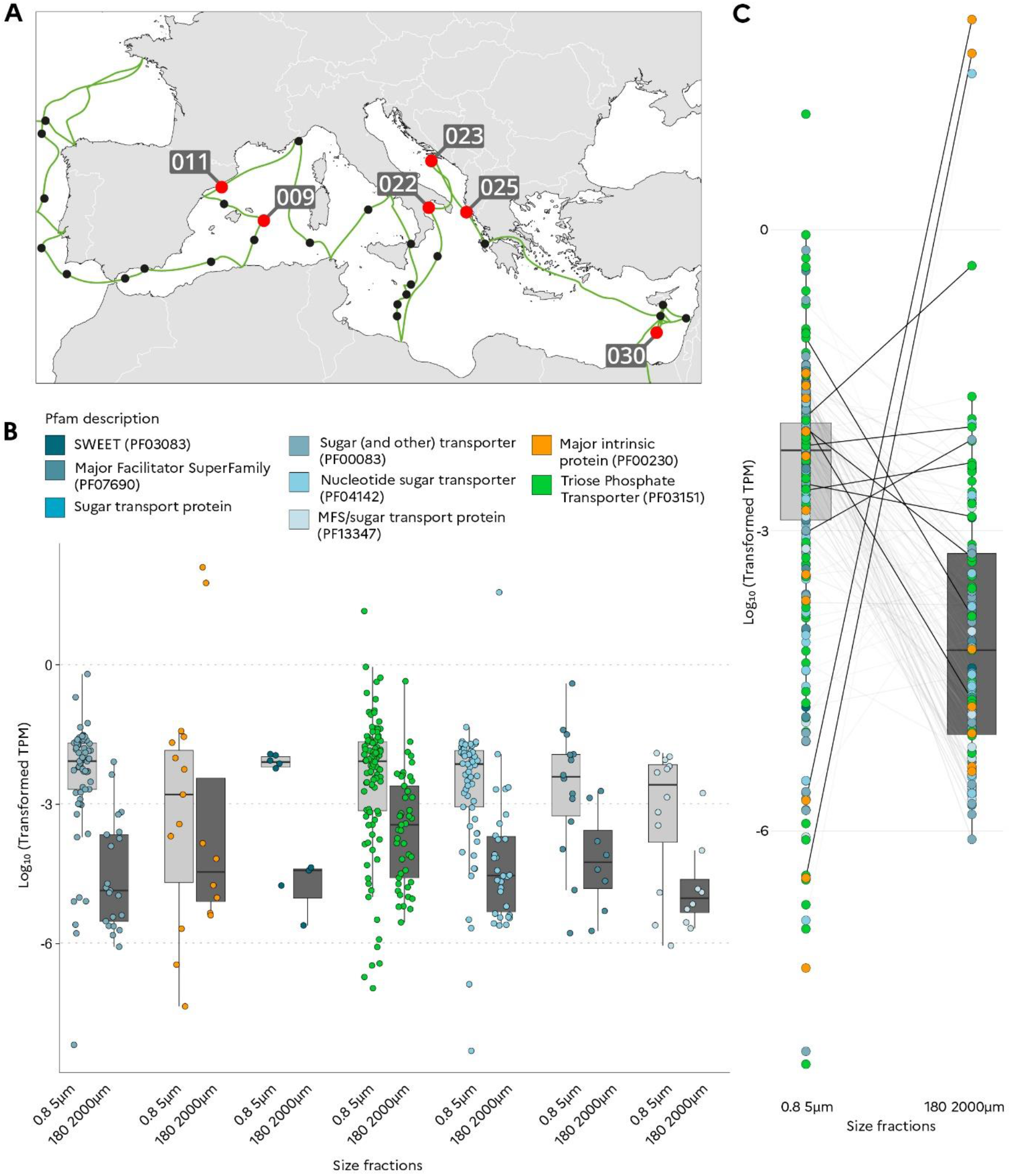
Normalized *in situ* expression of sugar transporter genes of the microalga *Phaeocystis cordata* in the Mediterranean Sea based on environmental metatranscriptomic data. **A)** Geographic map showing the six different sampling stations from the *Tara* Oceans expedition analyzed in this study. **B)** Comparison of the TPM (Transcript per Million) values for sugar transporter genes between the small (0.8-5 µm) and large (180-2000 µm) size fractions for each Pfam (each point corresponds to the TPM value of one sugar transporter gene). **C)** Comparison of TPM values of sugar transporter genes between the small and large size fractions including those upregulated in the single- holobiont transcriptomes (bold lines) (Fig. 3). In B and C, the TPM value of each sugar transporter gene was normalized by the expression of housekeeping genes (see Methods).

In order to normalize and compare gene expression between the two size fractions, we calculated the ratio between the expression (TPM) of the 270 sugar transporter genes and a selected set of 149 housekeeping genes (e.g. Ribosomal proteins, Tubulin, ATP synthase…) whose expression levels were expected to correspond to basic cellular activity (see Methods and Table S4). Based on this normalization method, we found 26 sugar transporter genes with a higher mean expression value in the large size fraction compared to the small one: twelve TPTs, six NSTs, three aquaporins, two Glycerol-Phosphate transporters, one GLUT, one SWEET and one member for the MFS/sugar transporter protein PF13347 (Fig. 4B, Table S4). Fourteen out of the nineteen sugar transporter genes found as upregulated in symbiosis (Fig.3B, single-holobiont transcriptomic analysis) were also found as expressed in both size fractions of the metatranscriptomic dataset. Of these, six of them presented a higher TPM value in the large size fraction: two aquaporin PF00230 genes, one of the two GLUT and three TPTs (Fig. 4C, Table S4). Note that the two aquaporin genes identified here correspond to the genes that presented the highest positive fold change values in symbiosis in the differential expression analysis from isolated holobionts (Fig. 3B). This *in situ* metatranscriptomic analysis provides an ecological significance to our experimental results, and further confirm some sugar transporter candidates as key players in symbiosis, such as TPTs, GLUT and aquaporin.

### Dynamic expression of the algal sugar transportome in symbiosis during the day

It is well established that photosynthesis and the central carbon metabolism depend on the circadian rhythm and light conditions [58–60]. Therefore, in order to further understand the metabolic connectivity, we investigated the transcriptional dynamics of the sugar transportome of symbiotic *Phaeocystis* by harvesting acantharian hosts (n = 4) at three different times of day and light exposure periods: 1) morning (9 am, after 1h light exposure), 2) evening (7 pm, after 10h light exposure) and 3) “dark-evening” (7pm, after an incubation in darkness for 24 hours). The “dark-evening” condition corresponds to a situation where holobionts, and thus microalgae experienced a non-photosynthetic day (absence of light). In total, we found 209 sugar transporter genes expressed in these three conditions, representing 96% (214/223) of the sugar transportome characterized above (Table S5, Table 1). Overall, 28% (60 genes) of sugar transporters were found to be expressed in the morning, 43% (91 genes) expressed in the evening and 29% (63 genes) in the “dark-evening” (Table S5). Among them, some were exclusively expressed in the morning (18), evening (43) or dark- evening (16) (Fig. 5A, Table S5). These results demonstrate that many sugar transporter genes of the symbiotic microalga tend to be induced during the day. From the comparison of holobionts collected in the evening and submitted or not to darkness (“dark-evening”), we found 53% (113/214) of genes downregulated when incubated in the dark (FC < −2, Table S5). 34% (72/214) were still expressed in the holobionts exposed to darkness and thus, their expression does not seem to be linked to the presence of light. These results show that the majority of sugar transporter expression of the symbiotic microalga is modulated by light conditions.

**Fig. 5.**
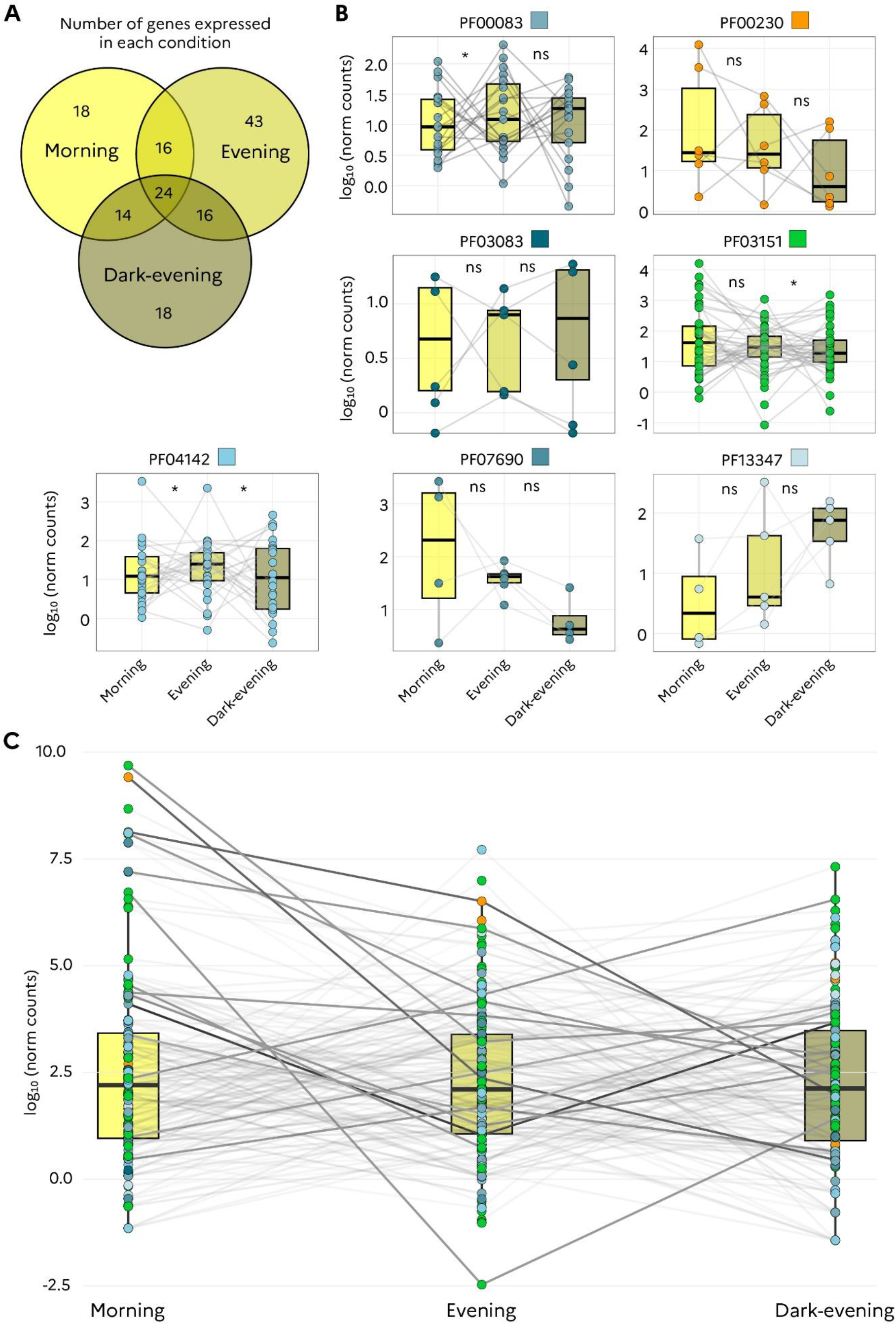
Temporal dynamics of the expression of the sugar transportome in the symbiotic microalga *Phaeocystis cordata*. **A)** Venn diagram showing the number of sugar transporter genes expressed in one or different conditions (morning, evening and dark-evening) (DESeq2 normalized read counts >10 for one condition and <10 for the other two conditions). **B)** Comparison of the normalized read counts of sugar transporter genes in different Pfams between morning, evening and dark-evening; ns (non-significant) and * (significant < 0.05) represent the Wilcoxon rank sum test significance for the comparisons between morning/evening and evening/dark-evening. **C)** Comparison of mean values of normalized read counts for the sugar transporter genes found upregulated in symbiosis (Fig.3B) across the three conditions (morning, evening and dark-evening).

At the Pfam level, we found specific expression patterns modulated by either light or period of the day (Fig. 5B, Table S5). For instance, the expression of TPTs and Glycerol 3-Phosphate transporter (PF07690) tended to be modulated only by light since they were significantly more expressed in evening vs “dark-evening” condition and did not show significantly higher expression between morning and evening (Wilcoxon rank sum test, *p*-value <0.05, Fig. 5B, Table S5). On the contrary, transcription of sugar hexose transporters (PF00083) seemed to be more regulated by the period of the day as shown by a significant higher expression in the evening compared to morning but not differentially expressed between evening and “dark- evening” conditions (Wilcoxon *p*-value<0.05, Fig. 5B, Table S5). Nucleotide sugar transporters (NSTs) seemed to be regulated by both parameters (light and period of the day) as they were found to be significantly more expressed in the evening *vs* morning, and “dark-evening” *vs* evening (Wilcoxon *p*-value <0.01, Fig. 5B, Table S5). MFS/sugar transport protein (PF13347) genes tended to present a higher expression in the dark (Fig. 5B, Table S5). SWEET genes did not present significant differences of expression levels between the three conditions, yet two pairs of genes seemed to present opposite patterns (more expressed in the morning or in the dark). Generally, aquaporin genes (PF00230) exhibited a lower expression in the darkness and two of them were only expressed in the evening, after the normal daylight exposure. These results show specific transcriptional patterns of sugar transporters according to each Pfam families responding to light (PF07690, PF03151, PF13347), to the period of the day (PF00083) or both parameters (PF04142).

We also paid attention to the dynamics of the sugar transporter genes found to be upregulated in symbiosis from our holobiont transcriptomes. The ten upregulated TPTs in symbiosis exhibited several transcriptional patterns: two genes exclusively expressed in the morning and two genes exclusively in the evening; in addition, two gene had a higher expression in the evening compare to morning and six a higher expression in light (evening) compare to dark-evening (Fig. 5C, Table S5). This suggests that different TPT genes might have specific roles at different periods of the day and this could depend on their subcellular localization. Most GLUT and aquaporin genes upregulated in symbiosis showed a higher expression in morning *vs* evening, or “dark-evening” *vs* evening, raising two hypotheses: 1) the transcription is activated in the morning to produce transporters during the day or 2) transcription mainly takes place in the dark for sugar excretion at night. Further studies should increase the temporal resolution during a day-night cycle to fully reveal the dynamics of the transportome expression of the symbiotic microalga.

## CONCLUSION AND PERSPECTIVES

This study improves our understanding of the molecular players that are potentially involved in the carbon metabolism and metabolic connectivity between the symbiotic microalga *Phaeocystis* and its acantharian host. We found that *Phaeocystis* species share a conserved sugar transportome among haptophytes at the Pfam level with few differences in gene copy number. Therefore, this genomic analysis did not reveal a specific sugar transportome linked to the symbiotic life stage of the microalga *Phaeocystis*, compared to non-symbiotic haptophytes. This can be explained by the fact that *Phaeocystis* symbionts are not vertically transmitted across host generations, do not depend on symbiosis for survival, and genome evolution would rather occur in the extensive free-living population [5]. Our study shows that the capacity of the microalga *Phaeocystis* to be in symbiosis may be rather due to the large plasticity of the transportome expression with 42% of transporter genes being differentially expressed. This suggests a drastic change in the flux and homeostasis of metabolites in the symbiotic microalgae. More specifically, downregulation of most transporter genes along with high expression of a few ones suggests i) lower trafficking of metabolites linked to the intracellular life stage (perhaps due to the arrested cell division) ii) specialization towards some metabolite fluxes putatively beneficial for the host. The transcriptional plasticity of the algal sugar transportome not only takes place during the free-living-to-symbiosis transition but also throughout the day. This reveals the complex dynamics of the carbon homeostasis and fluxes in the holobiont system.

Among the 19 sugar transporters of the microalga *Phaeocystis* upregulated in symbiosis, we found two GLUT and two aquaporin genes, which were also found more expressed in the large size fraction of environmental metatranscriptomics. This study provides further evidence that GLUT and aquaporin transporters, also found as upregulated in other photosymbiotic hosts [25, 26], play a key role in symbiosis. Similarly, the higher expression of a SWEET gene in this large size fraction suggests that this transporter may also be involved in metabolic connectivity in this planktonic symbiosis, as shown for anemone/dinoflagellate symbiosis [44].

The diversity of transporters and their expression patterns raise the hypothesis that several algal carbohydrates (glucose, glycerol) might be transferred to the host at different temporal windows. Future functional characterization (e.g. expression in heterologous systems) of the candidate sugar transporters revealed here will be essential to fully understand the role of these transporters in symbiosis. Overall, this study expands the list of holobionts using similar transporter genes and raises the hypothesis of a convergence for carbon exchange mechanisms in photosymbiosis.

## MATERIAL AND METHODS

### Dataset of genomic sequences of haptophytes transporters

Genomic identification of transporter genes was obtained from a re-annotation of the 186,115 protein sequences of six haptophyte species (*Diacronema lutheri, Emiliania huxleyi, Chrysochromulina tobinii, Phaeocystis antarctica, Phaeocystis globosa, Phaeocystis cordata*) from Phycocosm [61]. Sequences were re-annotated with multiple tools in order to have a complete description of each transporter useful for downstream analysis: Interproscan 5.60 with best scores for E-values < 0.000001 [62], TCDB Transporter Classification Database [63], blastp of the protein sequences with diamond (2.1.7, options: -e 0.00001 --ultra-sensitive --max-target-seqs 1) to the Uniprot release 2022 02, Hmmscan with Pfam-A.hmm 2021-11-15 [64], EggNog mapper [65]. To identify transporters, we first search in the merged files of annotations for the terms: “carrier|transport|channel|permease|symporter|exchanger|antiporter|periplasmic|facilitator”. We search sugar transporters using terms such as: “disaccahride|carbohydrate|sugar|ose|saccharide|glucose|polysaccharide”.

We used Phobius [66] and tmhmm 2.0 [67] to predict the number of transmembrane domains. We selected proteins that presented at least two transmembrane domains and less than three differences in terms of transmembrane domain number between Phobius and tmhmm. For PF00083 and PF07690 families, we used protein models (from InterPro database, https://www.ebi.ac.uk/interpro/) of the subfamilies domains described in Table S1, in order to build hmm profiles and use hmmsearch (best value, evalue E-23) to precise the identity of these transporters. Subcellular localization of *P. cordata* transporters was evaluated through five different tools: Deeploc 2.0 [68], TargetP 2.0 [69], Hectar 1.3 [70], WoLF PSORT [71], MuLocDeep 1 [72]. We used two thresholds 1) a Deeploc score >0.5 (as used in [73]) and 2) the consistency of prediction should be least the same for at least two tools, to select the most accurate putative predictions of transporters localization.

### Single-holobiont transcriptomics: sampling and analysis

For the free-living stage, a total of nine replicates of *Phaeocytis cordata* cells (strain RCC1383 from the Roscoff Culture Collection) maintained in K2 medium at 50-60 µmol PAR m^-2^s^-1^ and 20°C were harvested at 7pm at both late and stationary growth stages. Symbiotic acantharians with intra-cellular *Phaeocystis cordata* (holobionts) were collected with a 150µm plankton net in Mediterranean Sea in Villefranche-sur-Mer, France. Individual holobionts were manually isolated with a micropipette under a binocular microscope and rapidly transferred into filtered seawater (0.2 µm) and maintained in an incubator (50-75 μmol PAR m^-2^s^-1^, 20°C, 12h/12h). Free-living and symbiotic samples were frozen in the same conditions, in liquid nitrogen in a 0.2 µl PCR tube containing 4.4 µl of Smart-Seq2 buffer (Triton X-100 0.4 %/RNase inhibitor Ratio 19/1, dNTPs 10 mM, oligo dT 5 uM, [74]). Each sample was sequenced at 75 million reads, 2×150 paired-end with an Illumina NextSeq 500 instrument. A total of 1.9 billion reads were produced for this study.

Reads were first trimmed using trimmomatic (version 0.39, option PE -phred33; ILLUMINACLIP: contams_forward_rev.fa:2:30:10 LEADING:3 TRAILING:3 SLIDINGWINDOW:4:15 MINLEN:36 [75]) and bacterial, virus, human, fungi sequences contaminants were removed using kraken2 (2.1.2) and the k2_standard_202310 database [76]. In order to maximize the mapping rates of the reads, we built a new reference transcriptome of *Phaeocystis cordata* from the reads obtained with the sequencing of our culture (strain RCC1383 from the Roscoff Culture Collection) plus the reads from the PRJNA603434 BioProject deposited at NCBI GenBank [32] from the same *Phaeocystis* strain. Briefly, reads were assembled using rnaSPAdes v3.15.5 [77] and peptides were predicted using TransDecoder [78]. The peptides were annotated with the same method as for the genomic proteome. Conserved ortholog scores were calculated with BUSCO v5.4.4 [79] and alignment rates with Bowtie2 [80]. We reached a 68.87% re-mapping rate of the reads (31.13% with the previous reference, Figure S3); the two transcriptome references (this study and [32]) presented the same completeness (BUSCO score, Fig. S3 and [32]).

To verify if the decontamination of the reference transcriptome step using kraken2 was sufficient, we applied two blastp searches of the predicted peptides (as queries): 1) against *P. cordata* protein sequences from the Joint Genome Institute’s genome portal; and 2) against the NCBI nr database. For the 330 transcripts annotated as sugar transporter in the reference transcriptome, 96% of them were found in the protein sequences of *P. cordata* genome. The contigs presenting <50% identity (4 contigs) have a NCBI blast with *Oryza sativa*, *Arabidopsis thaliana* and 2 Bacteria but with a percentage of identity very close to the one found with the blastp against *P. cordata* genome-derived protein models. For the 14 proteins not found in *P. cordata* genome, only 4 have a NCBI match with a bacterial assignment but again with a <50% of identity. Thus, in total, six sugar transporters genes putatively presented a bacterial homolog but with ∼35% identity on average (Table S2, “merged_blastp”).

For the Differential Expression (DE) analysis, read counts were obtained using Kallisto (0.48.0) [81] to map the reads from holobionts and free-living RNA sequences on the reference transcriptome coding sequences. The differential gene expression analysis between free-living vs symbiotic stages was conducted using the DESeq2 R package (1.36.0, [82]). We used the threshold of normalized read counts >10 among all replicates to qualify a gene as expressed in a given condition [32].

### Tara Oceans metatranscriptomic data analysis

Reads of the Mediterranean stations 011, 009, 022, 023, 025 and 030 of the *Tara* Oceans expedition (2014) were obtained from the published dataset (PRJEB402/ERP006152, [55]). To compare the expression of sugar transporter gened between two size fractions, we transformed their TPM values into ratios between the expression of genes of interest and housekeeping genes that we identified in a similar way to the approach used by qPCR (Table S4 “HousekeepingGenes”). Briefly, the community expression patterns are compositional data, not absolute counts, and this type of data is constrained to an arbitrary fixed total defined by the sequencing depth, creating potentially spurious correlations through changes in abundance of other organisms in the system (see [83] for a review on the topic). A solution to this problem is to analyze ratios of gene expression instead of the proportions of the total. In this approach, choosing an appropriate denominator (housekeeping genes) to calculate ratios is critical [84]. We established a robust denominator via 4 criteria: (1) analyzing only samples in which at least 20% of the *Phaeocystis* transcriptome was expressed, and retain only genes expressed in all samples showing at least 20% *Phaeocystis* expression in every analyzed sample (this procedure eliminated 7 of the top 20 genes with the most reads mapped, likely because their high expression was the result of non-specific mapping in samples in which less than 20% of the *Phaeocystis* transcriptome was expressed), (2) selecting genes whose expression across all analyzed samples has a coefficient of variation below 200 (mean CV all genes= 374.8) and a fold change from the mean below 2, (3) checking that the genes present a functional annotation corresponding to typical housekeeping genes, and (4) checking that the genes correlate enough between them and through a k-means clustering, selecting the cluster with the highest amount of genes presenting r > 0.5 (adapting [84]). We then used the sugar transporters’ TPM as a numerator and the geometric mean of the housekeeping selected genes as a denominator to calculate our statistics.

### Temporal transcriptional dynamics of sugar transporters of the microalga *Phaeocystis cordata*

To evaluate the expression of the sugar transportome during the day, we collected more symbiotic acantharia in surface waters of the Mediterranean Sea (Villfranche- sur-Mer, France). For “Morning” samples, we sampled and isolated acantharia, maintained them in an incubator overnight (50-75 μmol PAR m^-2^s^-1^, 20°C, 12h/12h), and harvested them the day after at 9 am in the morning (one hour of light exposure). For “evening” samples, symbiotic acantharia were collected and isolated in filtered seawater, and frozen the same day around 7pm in the evening after 10 hours of light in the incubator. For “dark-evening” samples, holobionts were collected and maintained in the incubator with light until 8pm, and were transferred in a black box until 7pm of the following day. In order to compare gene expression across time/light conditions (morning, evening and dark-evening), we built a matrix of read counts for the three conditions using kallisto on the reference transcriptome of *P. cordata,* as explained above, and normalizing these counts using DESeq2 R package without “reference level” for the dds object creation.

## ACKNOWLEDGMENTS

Research was supported by CNRS and ATIP-Avenir program funding, and by the European Union (BIOcean5D - GA#101059915 and ERC – SymbiOCEAN -101088661). We thank the EMBRC program and the Institut de la mer de Villefranche-sur-Mer (IMEV, France) for the sampling. This project has received funding from the European Union’s Horizon 2020 research and innovation programme under grant agreement No 824110 for the sequencing (EASI Genomics). We thank Annelien Verfaillie and Alvaro Cortes from the Genomics Core plateform in Leuven. A. Auladell and D. J. Richter received funding from the European Research Council (ERC) under the European Union’s Horizon 2020 research and innovation program (grant agreement 949745 — GROWCEAN — ERC- 2020-STG) and from the Departament de Recerca i Universitats de la Generalitat de Catalunya (exp. 2021 SGR 00751). Z. Füssy and A. Allen were funded by National Oceanic and Atmospheric Administration grant NA19NOS4780181 (to AEA), the National Science Foundation (NSF-OCE-1756884 to AEA), and the Simons Collaboration on Principles of Microbial Ecosystems (PriME) (Grant ID: 970820 to AEA). The work (proposal: 10.46936/10.25585/60001426) conducted by the U.S. Department of Energy Joint Genome Institute (https://ror.org/04xm1d337), a DOE Office of Science User Facility, is supported by the Office of Science of the U.S. Department of Energy operated under Contract No. DE-AC02-05CH11231. We are grateful to the Roscoff Bioinformatics platform ABiMS (http://abims.sb-roscoff.fr), part of the Institut Français de Bioinformatique (ANR-11-INBS-0013) and BioGenouest network, for providing help for computing HECTAR subcellular localization prediction. We also thank Margaret Mars Brisbin for her help on the bioinformatic analyses.

## AUTHOR CONTRIBUTIONS

CJ and JD conceived, planned the research and wrote the manuscript; CJ, JD and FC performed sampling and experiments sampling; CJ performed data analysis, figures generation and drafted the first manuscript; AA, EP and DR performed data analysis from metatranscriptomics of *Tara* data. All authors participated in discussions interpreting results and provided comments and approved the final version.

## DATA AVAILABILITY

All raw sequencing data generated for this manuscript are available from the NCBI SRA under accession XXXXXXXXX.

