## Supplemental figures for "Transportome remodeling of a symbiotic microalga inside a planktonic host"

**This PDF file includes:**

Fig. S1 to S2

**Other supplementary materials for this manuscript include the following:**

Table S1. Excel

Table S2. Excel

Table S3. Excel

Table S4. Excel

Table S5. Excel

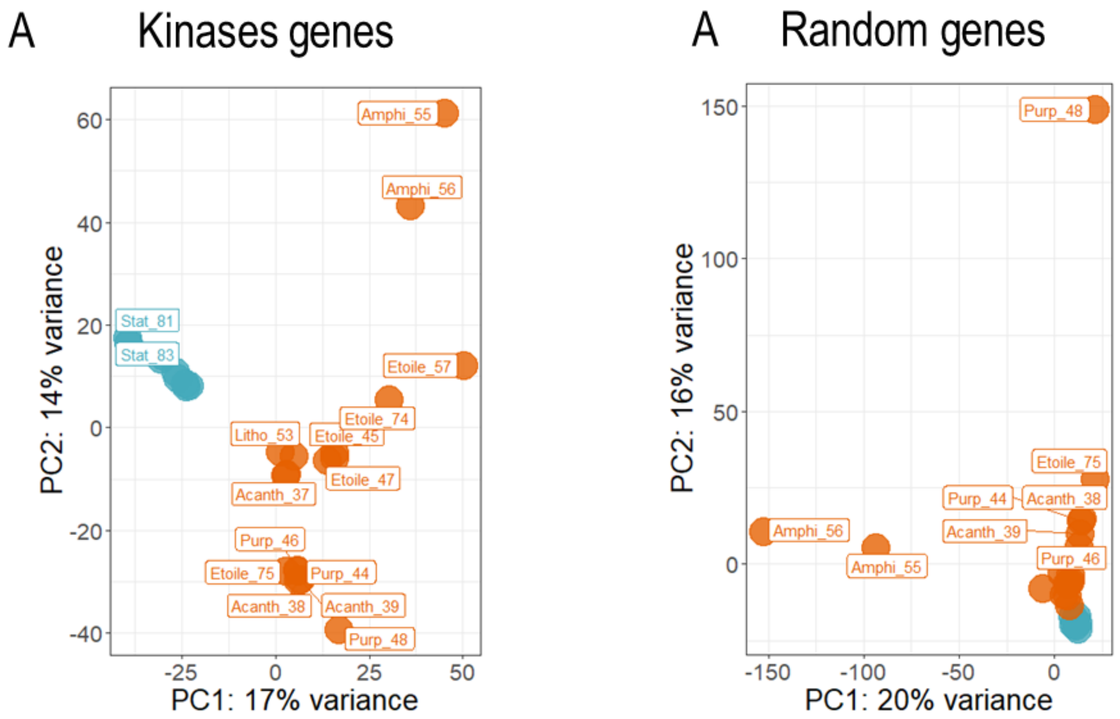


**Figure S1. Comparison of *Phaeocystis cordata* gene expression variance between Free Living and Symbiosis samples**. PCA of gene expression variance between conditions (x-axis) and between replicates (y-axis) for **A)** 1519 kinase genes and **A)** 2817 randomly selected genes.

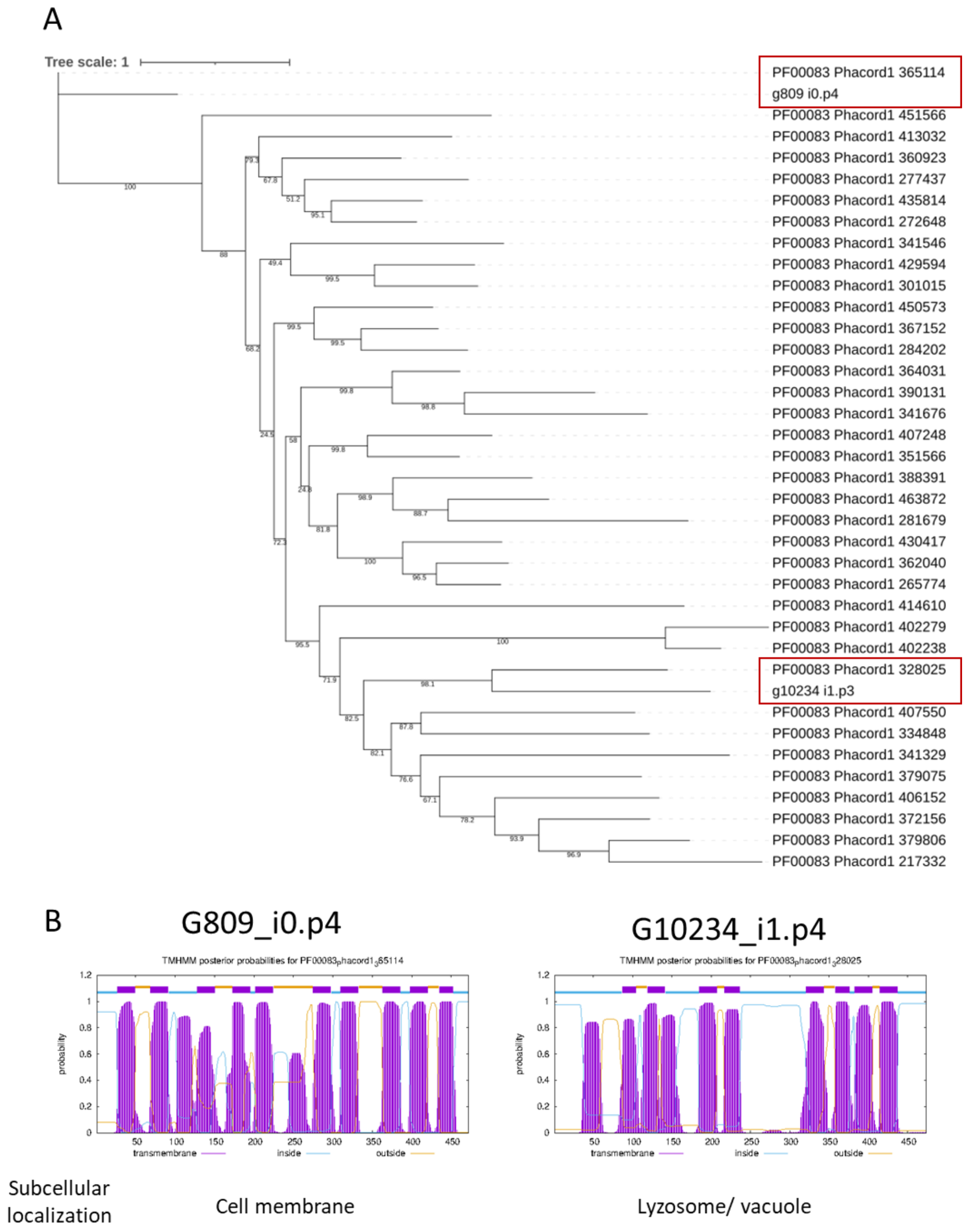


**Figure S2. A) Gene tree of PF00083 *Phaeocystis cordata* protein sequences from JGI and the predicted peptides from the reference transcriptome for the two upregulated genes in symbiosis.**

Model of substitution: Q.pfam+F+I+G4; 38 sequences with 1681 amino-acid sites; Number of constant sites: 575 (= 34.2058% of all sites); Number of invariant (constant or ambiguous constant) sites: 575 (= 34.2058% of all sites); Number of parsimony informative sites: 616. Number of distinct site patterns: 1329

**B) Transmembrane domain prediction of the two PF00083 predicted peptides upregulated in symbiosis** (TMHMM 2.0)

**A**

| | Mixed assembly Reads 1 + Reads 2 (spades) | Mixed assembly 1 and 2 CD-HIT-EST (spades) | Assembly Reads 1 (spades) | Assembly Reads 2 (spades) | Assembly reads from 2 (Trinity + CD-HIT-EST) | *P. Cordata* scaffold assembly |
|---|---|---|---|---|---|---|
| Nb transcripts | 112,171 | 104,281 | 137,446 | 58,110 | 32,621 | - |
| Nb peptides | 84,711 | 79,448 | 82,266 | 54,226 | 27,428 | - |
| Average lenght | 890 | 883.36 | 757 | 1124 | 1173 | - |
| busco | 67.8% (43,5% Single, 15% missing) | 67,8% (45,1% Single, 15% missing) | 58.8% (35,3% single, 25% missing) | 72% (50% Single, 14% missing) | 69% (12% missing) | - |
| Bowtie2 realignement (with the same sample) | 68.87% | 68.85% | 44.87% | 52.83% | 31,13% | 25.9% |
| Bowtie2 aligned concordantly exactly 1 time | 17,44% | 17.60% | 11.07% | 7,7% | 19,04% | 15.22% |
| Blastp on Phacord genomic proteome | 31,225 (87,6%) | | 29,606 (82,9%) | 31,196 (87,4%) | 25,995 (78,8%) | 35,688 |

**B**

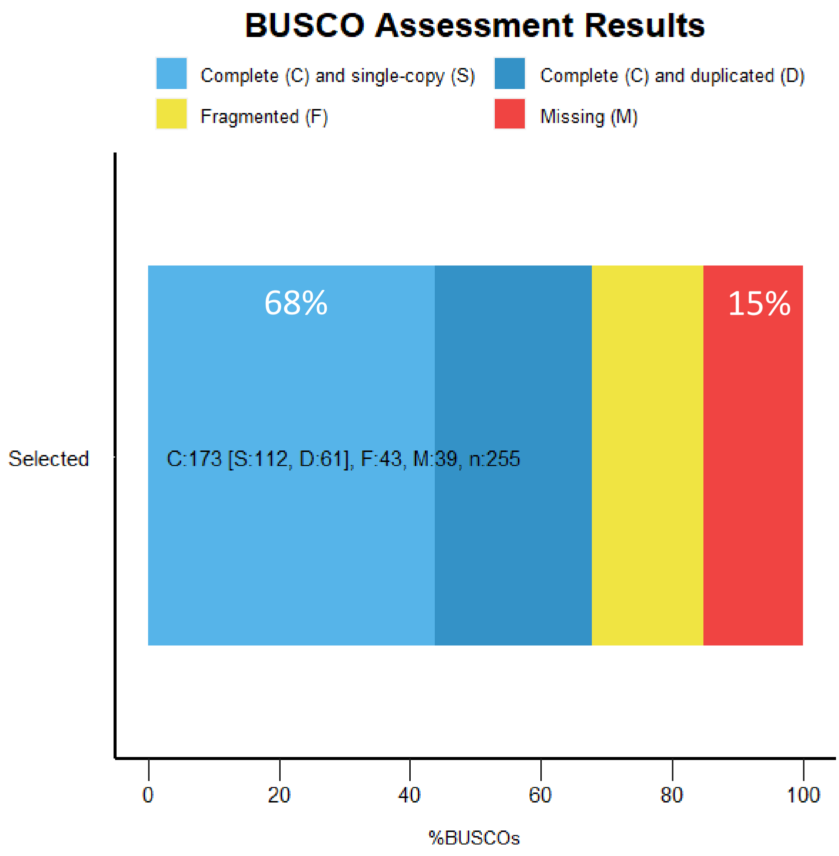


**Figure S3. Reference transcriptome statistics. A)** Comparison of multiple *Phaeocystis cordata* genome and reference transcriptome assemblies. Reads1 corresponds to the raw reads sequenced in this paper from the nine replicates of the culture of *Phaeocystis cordata;* Reads2 corresponds to the raw reads from PRJNA603434; *P. cordata* genome correspond to the JGI assembly. Two different tools were used for the reads assembly: Trinity V2.1.1 or spades V3.15.5. Re-alignment corresponds to the mapping of the reads used for the assembly on the references using bowtie2 V2.4.5

**B)** BUSCO score of the chosen reference transcriptome from A (Eukaryotic lineage).
